# Donor HLA-DQ reactive B cells clonally expand under chronic immunosuppression and include atypical CD21^low^CD27^−^ B cells with high-avidity germline B-cell receptors

**DOI:** 10.1101/2024.12.06.627284

**Authors:** Steven Russum, Ismail Sayin, Jasmine Shwetar, Eleonore Baughan, Jong Cheol Jeong, Andrea Kim, Alex Reyentovich, Nader Moazami, Adriana Zeevi, Anita S Chong, Marlena Habal

## Abstract

Long-term allograft survival is limited by humoral-associated chronic allograft rejection, suggesting inadequate constraint of humoral alloimmunity by contemporary immunosuppression. Heterogeneity in alloreactive B cells and the incomplete definition of which B cells participate in chronic rejection in immunosuppressed transplant recipients limits our ability to develop effective therapies. Using a double-fluorochrome single-HLA tetramer approach combined with single-cell *in vitro* culture, we investigated the B-cell receptor (BCR) repertoire characteristics, avidity, and phenotype of donor HLA-DQ reactive B cells in a transplant recipient with end-stage donor specific antibody (DSA)-associated cardiac allograft vasculopathy while receiving maintenance immunosuppression (tacrolimus, mycophenolate mofetil, prednisone). Donor DQB1*03:02/DQA1*03:01 (DQ8)-reactive IgG+ B cells were enriched for minimally mutated and germline encoded high avidity BCRs (median K_D_ 4.26×10^-09^) with an atypical, antigen-experienced and proliferative phenotype (CD27^-^CD21^low^CD71^+^CD11c^+/-^). These B cells coexisted with a smaller subset of more highly mutated, affinity matured IgG+CD27+ B cells. Circulating donor-reactive B cells and DSA remained detectable after rituximab, contrasting with the marked reduction in DSA after allograft explant and retransplant. Together, these findings define the persistence of germline high-avidity HLA-DQ alloreactive B cells and their co-existence with affinity matured clones that were both driven by the allograft despite conventional immunosuppression.

## 1. Introduction

Contemporary T-cell centric immunosuppression effectively controls early cellular rejection leading to excellent (>90%) 1-year survival in thoracic organ transplantation. However, it inadequately constrains humoral alloimmunity in both sensitized and non-sensitized recipients and fails to prevent chronic rejection^1–3^. Donor specific antibodies (DSA), predominantly directed against HLA class II antigens, develop in 25-30% of thoracic organ transplant recipients increasing the risk of both acute and chronic rejection that together portend a poor long-term prognosis^4–6^. Furthermore, in kidney transplantation, increased donor-reactive B-cell frequencies predict inferior outcomes in the absence of circulating DSA, which may be attributed to intragraft responses not detected in peripheral blood and/or to non-antibody mediated B-cell functions ^7^.

Despite the well described association between DSA and graft failure^4,8–11^, comparatively less is known about the cellular sources of DSA responses that develop and persist under contemporary immunosuppression. Following antigen encounter, B-cells are activated and upregulate surface HLA as well as costimulatory molecules that allow them to engage with cognate T-cell receptors (TCRs) on T-cells and provide reciprocal costimulation and cytokine signals required for B-cell survival^12^. The current paradigm of B-cell differentiation is that during the early response, B-cells differentiate into low affinity antibody secreting cells (ASCs) or early IgM+ memory B cells (Bmem). Activated B-cells that do not participate in this early response enter the germinal center and undergo affinity maturation: a tightly regulated process whereby coordinated and prolonged T-cell B-cell interactions drive the differentiation of ASCs and Bmem with progressively higher affinity as a result of somatic hypermutation (reviewed in ^13^). Notably, the mechanistic studies informing this paradigm underpinning our conceptual framework of humoral alloimmunity were performed in preclinical models in the absence of immunosuppression; how humoral alloimmunity evolves under immunosuppression that restricts T-cell responses in transplant recipients is not well understood.

More recent pre-clinical studies using lineage-tracking and alloreactivity models^14,15^ along with human B-cell analyses from individuals with autoimmune disease, suggest an alternative possibility; namely, high affinity B-cells can emerge from the bone marrow following stochastic BCR rearrangement and may have less stringent requirements for T-cell help. These findings therefore challenge the conventional paradigm that affinity maturation is a prerequisite for productive and persistent high avidity B-cell responses^14–18^.

The backbone of transplant immunosuppression relies on inhibiting T-cell activation through two main pathways; calcineurin inhibitors block signaling downstream of the T-cell receptor (signal 1) while costimulation blockade blocks the 2^nd^ signal required for effective T-cell activation. CNIs partially prevent the development of DSA; as evidenced by the higher incidence of DSA in CNI non-compliant transplant recipients and when CNI trough levels fall below 6ng/mL. However, DSA, in particular those directed against HLA-DQ antigens, develop and persist under calcineurin-inhibitor (CNI) based immunosuppression^9^ in up to 30% of transplant recipients. In contrast, costimulation blockade, which directly blocks T-cell/B-cell interactions, effectively prevents *de novo* DSA^19,20^, reverses ongoing humoral responses in preclinical models^21–23^ and suppresses anti-HLA antibodies in sensitized heart transplant candidates^24^, despite a higher incidence of T-cell mediated rejection. These observations suggest that incomplete attenuation of T-cell help by CNI-based immunosuppression permits de novo DSA responses and provides a potential explanation as to why they are controllable by costimulation blockade.

More recent pre-clinical studies using lineage-tracking and alloreactivity models^14,15^ along with human B-cell analyses from individuals with autoimmune disease, suggest an alternative possibility; namely, high affinity B-cells can emerge from the bone marrow following stochastic BCR rearrangement and may have less stringent requirements for T-cell help. These findings challenge the conventional paradigm that affinity maturation is a prerequisite for productive and persistent high avidity B-cell responses^14–18^, and further suggest the hypothesis that these stochastically generated high avidity B cells can differentiate into antibody secreting cells without entering into germinal centers, and are the major drivers of *de novo* DSA responses in immunosuppressed transplant recipients.

In this study, we sought to define the BCR repertoire characteristics, avidity, and phenotype of donor HLA-DQ reactive B cells that persist under conventional, CNI-based transplant immunosuppression. Using tools recently developed in the mouse model to specifically identify alloreactive B cells at single-cell resolution, we defined their phenotype, B-cell receptor (BCR) repertoire, and avidity in a case of DSA-associated end-stage chronic heart transplant rejection. From this single proof-of-principle case study, we provide evidence for the existence of high avidity donor HLA-DQ8 reactive B cells with germline-encoded BCRs that persisted under CNI-inclusive immunosuppression and in the absence of T-cell mediated rejection.

## 2. Methods

### 2.1 Sample processing and study approval

Peripheral blood was collected in heparin tubes under NYU Langone IRB #22-01226 in accordance with the declaration of Helsinki. Peripheral blood mononuclear cells (PBMCs) were obtained by Ficoll-density gradient, gradually brought to -80°C and transferred to liquid nitrogen for long-term storage. See supplementary methods and Table S1.

### 2.2 Multicolor flow cytometry and isolation of HLA tetramer binding B cells

PBMCs were stained with pre-titrated DQB1*03:02/DQA1*03:01 (DQ8) PE- and APC-conjugated tetramers followed by an 11-marker panel (each 30min, 4°C). Double-tetramer binding CD19+ B cells were index sorted (1 B cell/well) onto MS40L^lo^ feeder cells and cultured (see supplemental methods)^25,26^. Supernatants were tested for IgG and antigen specificity using pooled 5-peak carboxyl beads conjugated to biotinylated antigen monomers, stained with PE-conjugated anti-IgG and analyzed (BioRadZE5).

### 2.3 B-cell receptor sequencing and recombinant antibody production

Expanded B-cell clones (2.2 above) were snap frozen in RLT with 1% mercaptoethanol then used for BCR sequencing (iRepertoire, Huntsville, AL). Sequences were annotated and analyzed using IMGT/V-Quest and NCBI Blast. Clones were defined as having the same V_H_, J_H_, identical CDR3 length, and >85% CDR3 sequence homology. Heavy and light chain inferred germline sequences were reconstructed (Change-O) after aligning to the reference sequence (IMGT) and mutated residues reverted to the reference sequence without modifying non-templated nucleotide additions. Vectors carrying the heavy and light chains were co-transfected into HEK293 cells, cultured, and supernatants extracted for affinity purification (Sino Biological).

### 2.4 Affinity determination by serial dilution and surface plasmon resonance (SPR)

IgG concentration in the culture supernatants was determined from a standard curve and the relative avidity of each clone was determined by serial titration. The K_D_ by limiting dilution was determined using a nonlinear regression 1-site binding model. Avidity by SPR (GE Biacore 8K) was determined using the Series S sensor chip (Cytivia) and immobilized biotinylated DQB1*03:02/DQA1*03:01 (25nM). Serially diluted supernatants and recombinant monoclonal antibodies (range 20nM to 1nM) were run over the sensor chip for 120s followed by a dissociation step (240-600s). K_D_ (SPR) was determined using 2-state reaction kinetics (Biacore analysis software).

## 3. Results

### 3.1 Expanded switched, IgG+ donor-reactive B cells from a transplant recipient with end-stage cardiac allograft vasculopathy predominantly express germline or minimally mutated B-cell receptors with high-avidity for donor HLA-DQ8

In transplant recipients with chronic rejection, persistent DSA is associated with inferior outcomes despite maintenance immunosuppression. To define the donor-reactive B cells that expanded in the setting of immunosuppression (tacrolimus (TAC), mycophenolate mofetil (MMF), Prednisone), PBMCs were obtained from a transplant recipient with end-stage cardiac allograft vasculopathy (CAV) and circulating HLA-DQ8 DSA at the time of a biopsy without evidence of acute rejection (ACR grade 0, AMR grade 0). Using a combination of PE- and APC-labelled HLA DQB1*03:02/DQA1*03:01 (hereafter referred to as HLA-DQ8) tetramers that had been titrated to reduce background, the frequency of donor HLA-DQ8 reactive B cells was 0.64% within switched IgG^+^ B cells (Fig 1A). In contrast the frequency of IgG+ HLA-DQ8 reactive B cells in the control was <0.05%.

**Figure 1.**
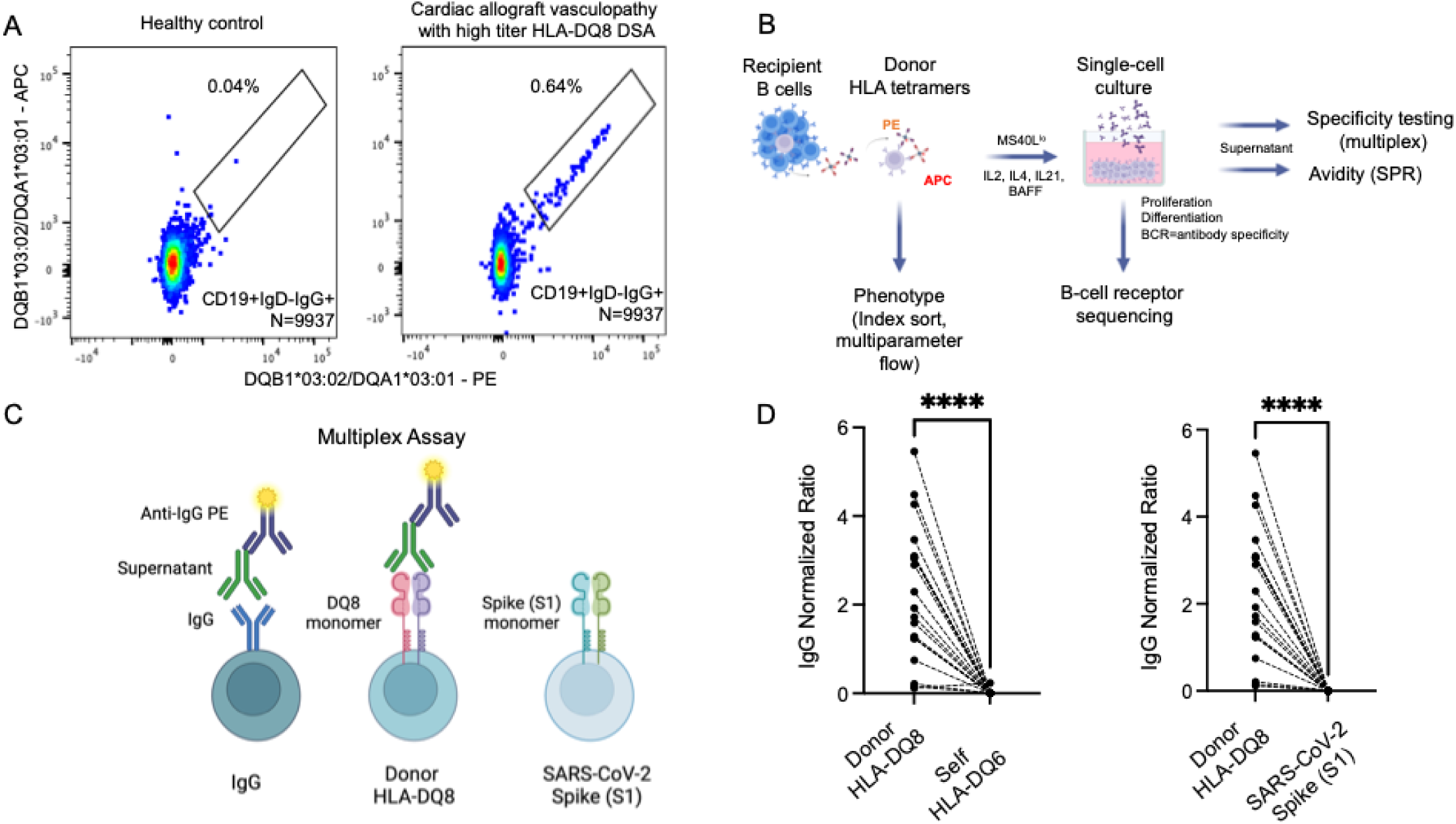
HLA tetramers specifically identify IgG+ donor-reactive B cells. (A) Fluorochrome labeled HLA-DQ8 tetramers were titrated to detect donor-reactive IgG^+^ B cells in peripheral blood while reducing non-specific binding of control B cells. (B) Experimental approach to sorting HLA-DQ8-binding B cells, *in vitro* differentiation into antibody secreting cells, testing the specificity of secreted IgG and its avidity for donor HLA-DQ8. (C) IgG coated beads were used to quantify total IgG in the culture supernatant, and the antigen-specificity of the IgG was assessed with beads coated with donor HLA-DQ8 or irrelevant antigen (SARS-CoV-2 Spike S1). Results are presented in (D). **** p<0.0001, paired t-test. Fig B, C created with biorender.com.

To confirm BCR specificity, tetramer-binding B cells were index sorted (1 B cell/well) and cultured with CD40L-expressing stromal cells (MS40L^low^) as previously described^25,26^ (Fig 1B). Single-B cells proliferated and differentiated into antibody-secreting cells (ASCs), thus allowing the specificity of their secreted IgG to be assessed using single-antigen beads coated with the same donor DQB1*03:02/DQA1*03:01 HLA. Specificity controls included self-HLA or an irrelevant antigen (SARS-CoV-2 spike protein (S1); Fig 1C). Cloning efficiency for the IgG+ sorted donor-reactive B cells was 79% (19/24) and all clones tested bound to HLA-DQ8 but not to self-HLA or S1 (Fig 1D). Hence, the tetramer-binding assay was highly specific in identifying IgG+ HLA-reactive B cells in this transplant recipient.

These studies demonstrated that IgG+ donor HLA-DQ8 reactive B cells persisted at high frequencies in this patient with CAV and circulating HLA-DQ8 DSA, even though alloreactive effector T-cells were absent from the allograft (TCMR grade 0). We proposed two non-mutually exclusive possibilities to explain the persistence of these B cells in the setting of limited effector T-cell responses. First, the HLA-DQ8 reactive B cells could be highly mutated, affinity matured ‘classical Bmem’ that developed remotely during brief periods of reduced immunosuppression and are persisting in a quiescent state. Alternatively, they could be minimally mutated clones that are actively responding to the allograft despite high-dose maintenance immunosuppression (TAC with trough levels >6ng/ml, MMF 1500mg BID, Prednisone 10mg/d). Understanding how these distinct B cell differentiation pathways may contribute to the chronic humoral response under immunosuppression would be informative to guide the development of more effective therapies in the clinic.

To address this question, we first sought to define the BCR heavy and light chain sequences of the identified HLA-DQ8 reactive IgG+ B cells. Although there was heterogeneity in somatic hypermutation frequency, most (68.4%) BCR IGHV sequences had fewer than 7 mutations (mutation frequency <2.5%) and strikingly three BCRs had a germline configuration (no mutations) when referenced to the IMGT database (Fig 2A,B). In contrast to the B cells with a low mutational burden, the more mutated clones, had a higher ratio of non-silent to silent mutations possibly suggestive of affinity selection (Fig 2C, Table S2). Notably, amongst the 19-donor specific IgG+ B cells that were sequenced, two expanded clonotypes were identified (defined as sharing the same V_H_, J_H_, and CDR3 length with >85% CDR3 homology, and the same light chain; Fig 1D). One was a minimally mutated clonotype that represented four of the nineteen (21%) donor specific IgG+ B cells sequenced (IGHV nucleotide (NT) mutations, 5,5,5,1; clone 1). The second clonotype of 2 clones had a higher mutational burden (NT mutations 17 and 13; clone 2). Collectively these findings demonstrate the expansion and coexistence of alloreactive B-cells with high, low or no mutations and suggest that affinity maturation is not a prerequisite for donor alloreactivity in immunosuppressed transplant recipients.

**Figure 2.**
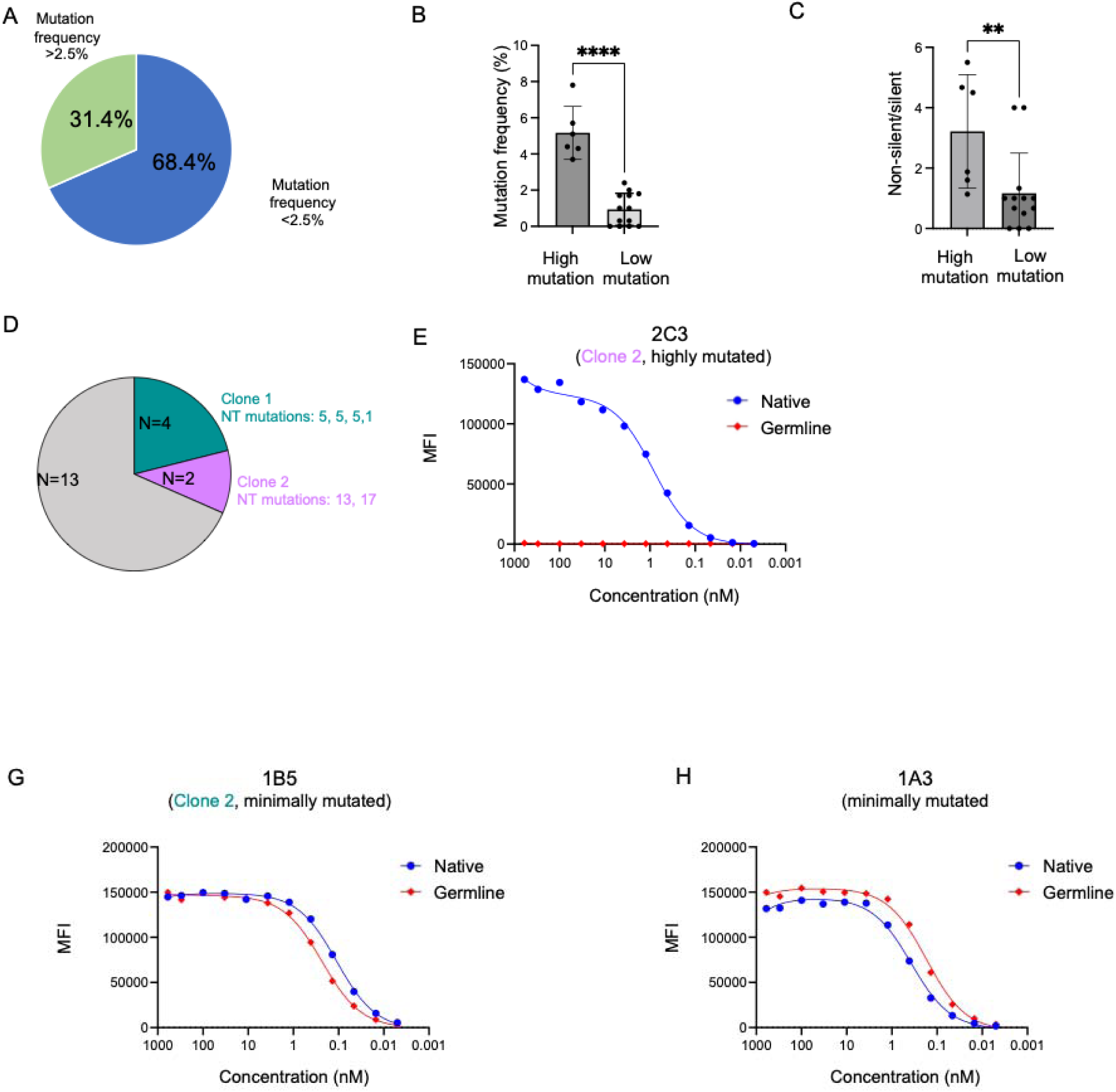
Repertoire characteristics of IgG+ donor HLA-DQ8 reactive B cells. (A) Percentage of highly mutated (mutation frequency >2.5%) versus germline and minimally mutated (mutation frequency <2.5%) donor reactive B cells, (B) Mutation frequency within each group (high vs. low mutation frequency), (C) ratio of non-silent to silent mutations, (D) expanded clones within IgG+ donor reactive B cells, (E-G) rAb titration for the native BCR compared to its unmutated inferred germline for 3 different B cells. (E) Affinity maturation of the highly mutated BCR (2C3, clone 2) for donor HLA-DQ8 compared to the inferred germline. (F,G) Both native and inferred germline rAb for the minimally mutated BCRs (1B5, clone 1 and 1A3) had similar binding to donor HLA-DQ8. NT, nucleotide; Nb, number; rAb, recombinant monoclonal antibody. ** p<0.01, paired t-test; box plot error bars (Fig B and C) show minimum and maximum values.

Having identified enrichment for minimally mutated alloreactive B cells, we wanted to rigorously test if they were high or low-avidity B-cells. The normalized MFI ratio (donor-HLA/total IgG) used to define antigen specificity suggested that most of these clones were high or mid-avidity B cells (mean ratio 2.4; range 0.15-5.5), and we further confirmed this conclusion using dilution series as well as surface plasmon resonance (SPR), a gold standard assessment of binding avidity. The generated Kd curves using MFIs (representative in Fig S1) and SPR confirmed that donor-reactive IgG+ B cells, including germline and minimally mutated BCRs, had very high affinity for donor HLA-DQ8 (median K_D_ 4.13×10^-09^).

Two potential scenarios could explain the presence of minimally mutated, high avidity clones. One was that the BCRs had undergone strong selection through classical affinity maturation resulting from mutations at a few critical residues. Alternatively, it was possible that these B cells had germline high affinity BCRs and that the mutations had minimal impact on their affinity. To address these possibilities, we first inferred, unmutated germline sequences for three B-cells (IMGT/HighV-QUEST) and then expressed them as recombinant monoclonal IgG (rAbs); two were minimally mutated (1B5 from clone 1, and 1A3) and one was highly mutated (2C3 from clone 2). For each clone, we generated a pair of rAbs derived from the sequenced BCR (‘native’) and inferred unmutated sequence.

For the highly mutated B cell (2C3 from clone 2), the germline rAb had significantly reduced donor HLA-DQ8 binding compared to the native rAb (Fig 2E). This cloned 2C3 B cell had intermediate-high avidity K_D_ (29.8nM) while the germline revertant did not appreciably bind to HLA-DQ8 by SPR (Table 1, Fig S2). Thus, this is a B cell clone that had undergone affinity maturation. In striking contrast, for both the minimally mutated B cells (1B5, clone 1 and 1A3), the germline reverted rAb maintained comparable binding to donor HLA-DQ8 as their respective native rAb (Fig 2F,G). Both native minimally mutated BCRs had high avidity (2.6nM, 5.8nM) for donor HLA-DQ8, and this avidity persisted (1.7nM, 4.1nM) in the germline reverted rAbs (Table 1, Fig S2). Taken together these studies provide strong evidence for the co-existence of high avidity germline and somatically mutated, affinity matured clones in this one case of chronic rejection despite conventional immunosuppression.

**Table 1.** K_D_ of recombinant monoclonal antibodies from alloreactive B-cell receptor sequences and their predicted unmutated germline sequence.

| **Recombinant**<br>**(native)\*** | **$K_D$ (nM)** | **Recombinant**<br>**(germline,**<br>**unmutated)\*\*** | **$K_D$ (nM)** |
| --- | --- | --- | --- |
| **2C3 native** | 29.8 | **2C3 germline** | No appreciable<br>binding |
| **1B5 native** | 5.8 | **1B5 germline** | 4.1 |
| **1A3 native** | 2.6 | **1A3 germline** | 1.7 |

\*Native denotes the B-cell derived heavy/light chain sequence.
\*\*Germline denotes the reverted, unmutated germline predicted sequence.

### 3.2 High avidity IgG+ donor HLA-DQ8 reactive B cells are enriched for an antigen-experienced, atypical memory phenotype

To further characterize the donor-reactive B-cell response in this immunosuppressed transplant recipient, we phenotyped the index-sorted donor-reactive B-cells using a panel of markers that are associated with antigen experience, memory, and activation. Unsupervised clustering analysis showed that donor-reactive switched B cells had a distinct phenotype relative to total switched B cells (Fig 3A) characterized by enrichment within the CD27^−^IgD^−^ ‘double negative’ (DN) subset (Fig 3B). These donor-reactive B cells showed a preponderance towards antigen experience (CD21^low^) and an atypical (CD19^high^,CD11c^+^) phenotype previously described as extrafollicular (EF) differentiated B cells in autoimmunity and infection^17,18^ (Fig 3C). Donor-reactive B cells also expressed higher levels of CD71, consistent with inadequate constraint of proliferation. Together, enrichment for a minimally mutated BCR repertoire and an atypical phenotype suggest that many of these B-cells may have developed through a non-classical ‘atypical’ pathway defined as not requiring stringent GC-associated T-cell help and affinity maturation. These observations suggest a novel potential mechanism for the humoral response that developed in this compliant transplant recipient who was taking high-dose immunosuppression.

**Figure 3.**
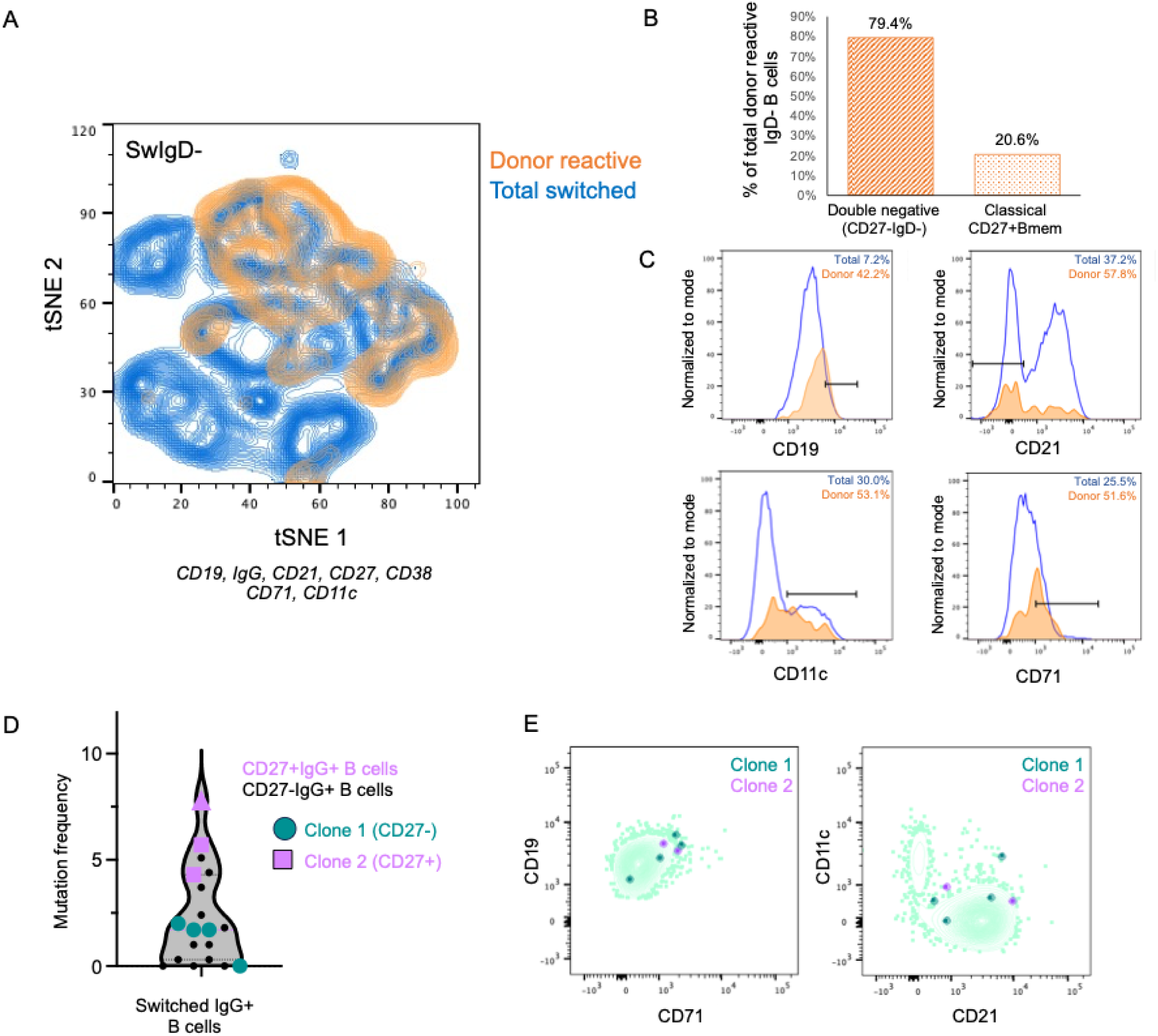
Donor specific IgG+ B cells are enriched for a CD27^-^CD21^low^ antigen experienced phenotype. Unsupervised cluster analysis (A) of total switched (SwIgD^-^) B cells (blue) to demonstrate that donor-reactive SwIgD^-^ B cells (orange) are enriched within the CD27^-^ subset (B), and they express markers associated with antigen experience (C; blue line, total CD19^+^IgD^-^IgG^+^ B cells; orange, donor-reactive CD19^+^IgD^-^IgG^+^ B cells), CD27+ donor reactive B cells have mutated BCRs (D), expanded clones include cells with a phenotype of antigen-experience, atypical memory (CD21^low^, CD19^high^, CD11c^+/-^) and proliferation (CD71^int/high^). MFI, mean fluorescence intensity.

In contrast to atypical B cells, classical Bmem that undergo affinity maturation are frequently identified by the canonical memory marker CD27. Hence, we determined whether the B-cells with the highest mutation frequencies were more likely to be CD27+. Consistent with this paradigm, highly mutated donor reactive IgG+ B cells were CD27+ (Fig 3D); including the affinity matured expanded clone (clone 2, purple squares in Fig 3D). Conversely, Clone 1 which had minimally mutated BCRs that maintained donor specificity in the germline reverted state was part of the CD27-subset (green circles in Fig 3D).

To address whether donor-reactive B cells expanded remotely leaving behind quiescent circulating clones or whether they were the product of an ongoing response, we focused on markers of proliferation (CD71) and antigen experience (CD21^low/int^ and/or CD11c^hi^). Superimposing clones 1 and 2 onto the transplant recipient’s global B-cell phenotype showed that 5 of the 6 B cells in the two expanded clones were CD71^int/high^ and 3 of 6 clones had downregulated CD21 with variable CD11c expression (Fig 3E). These results are consistent with the clonally expanded B cells from both classical Bmem and atypical subsets being in an active state.

### 3.3 High avidity donor reactive pre-switched, IgD+ B cells with germline B-cell receptors are also detectable within the circulation and harbor an antigen experienced phenotype

The unexpected observation that switched, IgG+ B cells maintained high avidity for donor HLA when their BCR was reverted to the germline, prompted us to ask whether ‘naïve-like’ pre-switched (IgD+) B cells with donor-reactive germline BCRs might also be present in the circulation and if so, whether these B cells were antigen experienced. Hence, we single cell sorted CD19+IgD+ tetramer binding B cells into Nojima cultures and screened the supernatants to identify donor-reactivity using a conservative MFI cut-off of 1000 (n=21, Fig 4A). Index-sort phenotype demonstrated that 20/21 of the pre-switched donor HLA-DQ8 -reactive B cells did not express the canonical memory marker CD27. Nonetheless, most donor reactive pre-switched B cells were found within the CD21^low^ subset as defined by total double-tetramer binding B cells (Fig 4B, S3) and within the antigen verified clones (Fig 4C) suggesting that these B cells were recently activated and antigen experience. We next selected 8 pre-switched IgD+ clones for BCR sequencing. These studies confirmed that donor-reactive IgD+ B cells expressed germline BCR sequences (≤ 1 mutation) and identified an expanded, unmutated IgD^+^CD21^-^ clone (n=2). Hence, our finding demonstrate that alloreactive B-cell activation and clonal expansion can occur before isotype switching.

**Figure 4.**
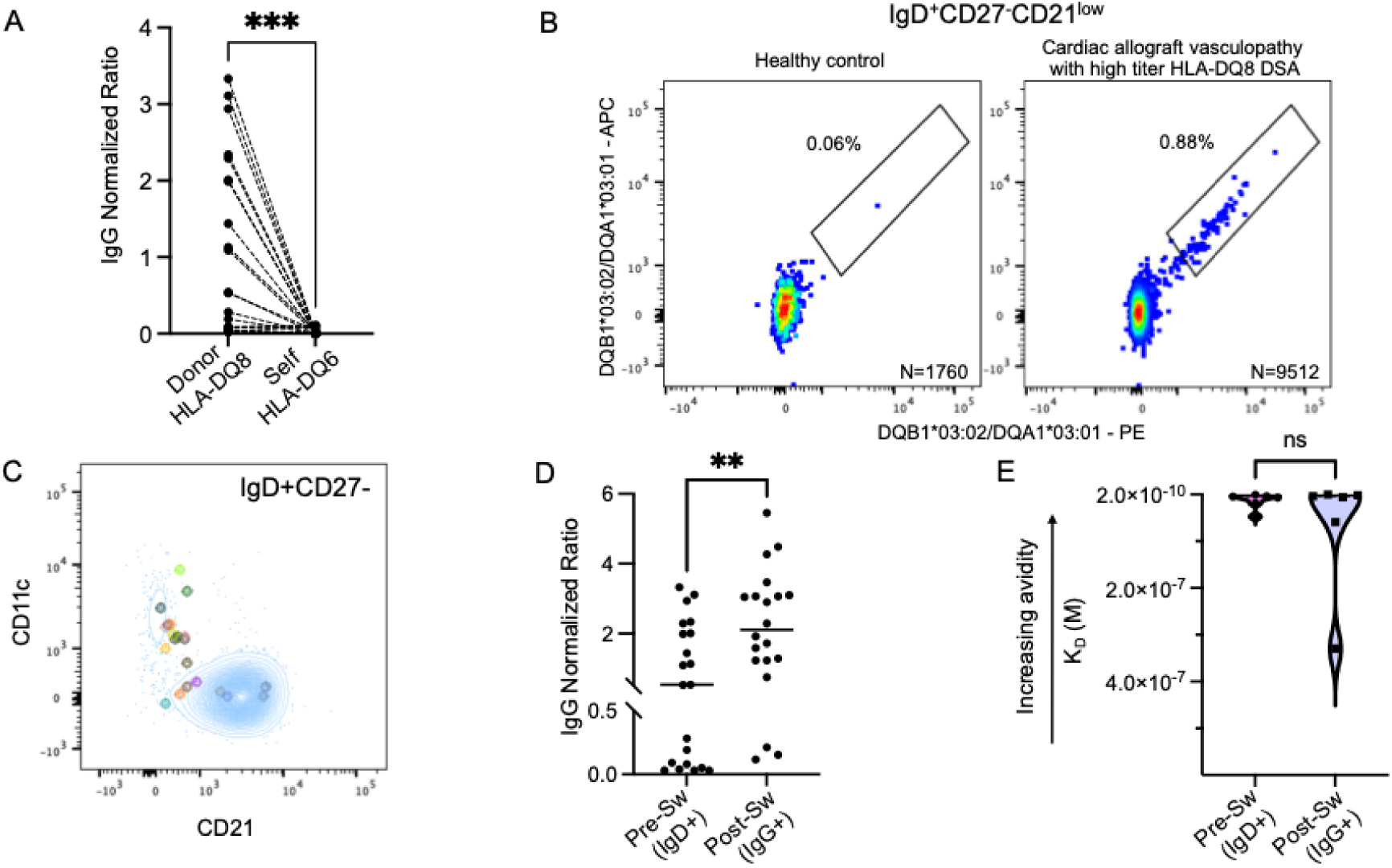
Unmutated pre-switched IgD+ donor-reactive B cells include high avidity clones. (A) Normalized IgG MFIs of single B-cell culture supernatants; HLA-DQ8 binding and pre-switched B cells were *in vitro* differentiated into isotype switched IgG antibody secreting cells as described in Fig 1. Positive supernatants were defined by an MFI>1000 on HLA-DQ8 beads. (B) Enrichment of donor-reactive B cells in the pre-switched IgD^+^CD27^-^CD21^low^ B-cell subset from peripheral blood of transplant recipient compared to control. (C) Verified donor-reactive preSw B cells superimposed onto the transplant recipient’s global pre-switched (IgD^+^CD27^-^) B-cells confirming enrichment for CD21^low^, antigen experienced phenotype and variable expression of CD11c. (D) IgG normalized MFI of culture supernatants from pre-switched (IgD+) vs. post-switched (IgG+) donor HLA-DQ8 reactive B-cell clones; (E) SPR defined K_D_ of a subset (see Fig S3) of pre-switched B cells with high normalized IgG ratios compared to IgG+ clones, PreSw, pre-switched IgD^+^CD27^-^; Bmem, memory B cells; DN, double negative. *** p<0.001, **p<0.01; paired t-test (in A), unpaired t-test (in D and E).

We next sought to determine whether these pre-switched IgD+ donor reactive B cells might also have high avidity for donor HLA alloantigens analogous to the germline revertants. We first used the normalized HLA-DQ8 MFI as a surrogate for avidity and found that in addition to some lower reactivity clones, a substantial number of donor-reactive IgD+ B cells had a similar normalized MFI to the IgG+ B cells (Fig 4D). We next confirmed by SPR that this subset of pre-switched B cells had high avidity for donor HLA (K_D_ range 6.2×10^-11^M to 4.8×10^-8^M) that was not significantly different from the K_D_ ranges of switched IgG+ B cells (K_D_ range 4.9×10^-10^M to 3.3×10^-7^M; Fig 4E, Fig S4, Table 2). Collectively, the detection of high avidity donor-HLA-DQ8-reactive germline BCRs in both the pre-switched and post-switched compartment along with the finding that germline revertants maintained high-avidity for donor HLA DQ8 confirmed that affinity maturation was not required in order to develop a clonally expanded, high-avidity donor-reactive B cell repertoire in this transplant recipient with end-stage CAV.

**Table 2.** K_D_ of pre-switched IgD+ and post-switched IgG+ alloreactive B-cells used in the SPR analysis.

| **Clone** | **Phenotype** | **$K_D$\*** |
| --- | --- | --- |
| 2E1 | IgD+CD27- | 4.75E-08 |
| 1F10 | IgD+CD27- | 1.89E-08 |
| 2B4 | IgD+CD27- | 6.98E-09 |
| 1E5 | IgD+CD27- | 4.64E-09 |
| 2F11 | IgD+CD27- | 3.87E-09 |
| 1H2 | IgD+CD27- | 6.24E-11 |
| 1G7 | IgD+CD27+ | 6.57E-10 |
| 2A7 | IgG+CD27- | 3.30E-07 |
| 1H12 | IgG+CD27- | 5.92E-08 |
| 1F5 | IgG+CD27- | 5.67E-09 |
| 1G12 | IgG+CD27- | 3.76E-09 |
| 1B5 | IgG+CD27- | 4.94E-10 |
| 2A9 | IgG+CD27+ | 1.84E-09 |

\*Studies were performed using supernatants from the single B-cell cultures.

### 3.4 Rituximab does not adequately deplete donor-reactive B cells

Monotherapy with rituximab or proteasome inhibitors has not been successful in reversing chronic rejection^27–29^. Although there are various potential explanations, one possibility is that *de novo* differentiation of antigen specific B-cells from naïve and/or Bmem repertoires continually replace antibody-secreting cells, and that this process is potentiated by increased BAFF after B-cell depletion^30^. To investigate this possibility, we studied B cells after the abovementioned transplant recipient had been clinically treated with rituximab and carfilzomib. Three days after the first treatment with rituximab, a substantially reduced frequency of circulating CD20^-^CD19^dim^ B-cell population was detected (Fig 5A); however, within this subset we were able to isolate, sort, and single-cell culture donor HLA-DQ8 tetramer binding B cells (Fig 5B,C). These B cells harbored a CD21^low^CD71^int^ phenotype consistent with recent activation and proliferation (Fig S5). One month later, after another dose of rituximab and the addition of carfilzomib, we were still able to detect donor-reactive B cells based on HLA-DQ8 tetramer binding (Fig 5D) although we were unsuccessful in culturing these B cells possibly due to exposure to the irreversible proteasome inhibitor carfilzomib. Hence, our findings provide an explanation for the limited efficacy of current therapeutic strategies that do not adequately constrain B-cell differentiation.

**Figure 5.**
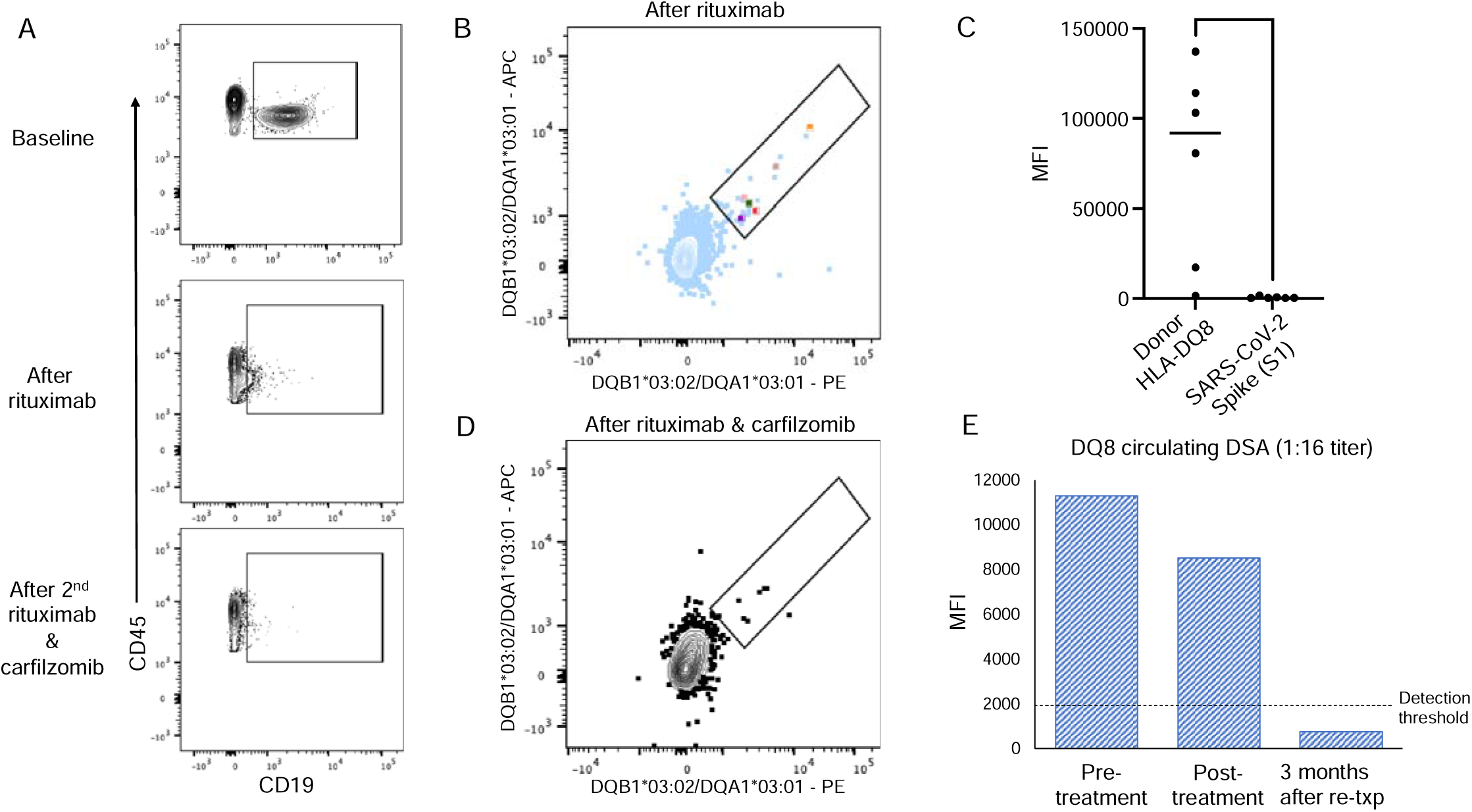
Humoral response after rituximab and carfilzomib or removal of the cardiac allograft. B cells (live gated CD45^+^CD3^-^CD14^-^) before and after rituximab and carfilzomib treatment. (B) Donor-reactive B cells detected after anti-CD20 therapy (rituximab). Colored dots represent single-cell CD19^dim^CD20^-^ B cells that were cultured and confirmed for donor specificity. (C) Supernatants of B-cell clones were tested for binding to HLA-DQ8 (donor) vs. SARS-CoV-2 S1. (D) Donor-reactive B cells detected after rituximab and carfilzomib. (E) Serum HLA-DQ8 DSA before and after rituximab and carfilzomib treatment and 3 months after allograft explant and retransplant with an allograft that was HLA-DQ matched to the recipient. MFI, mean fluorescence intensity; Txp, transplant; DSA, donor specific antibodies; *p<0.05, paired t-test.

Although rituximab and carfilzomib treatment was ineffective in reducing circulating DSA, three months after explant of the allograft and re-transplant with a heart that was HLA-DQ matched to the recipient, we observed a profound decrease in circulating DSA (Fig 5E). This suggested that antibody persistence in the presence of an allograft can be reversed once the antigenic stimulus is removed. These findings provide additional support for a dynamic response in which ongoing B-cell priming by donor antigen results in their differentiation into short-lived antibody-secreting cells that actively contribute to DSA persistence.

## 4. Discussion

In this study, we identified high avidity donor HLA-DQ reactive B cells with BCRs that maintained a germline or minimally mutated configuration in a single-case of end-stage CAV with high titer HLA-DQ8 DSA. This proof-of-concept observation challenges the conventional paradigm that was built on preclinical models wherein immune responses were investigated without immunomodulation by therapies widely utilized in transplant recipients. In those studies, post-germinal center B cells with evidence of affinity maturation are a dominant feature of pathogenic anti-donor-HLA responses. The striking enrichment for CD27-B cells expressing an antigen experienced and proliferative phenotype across the pre- and post-switched subsets in this transplant recipient attests to their prevalence and resistance to both anti-metabolite (MMF) and T-cell centric calcineurin inhibitor (tacrolimus)-based immunosuppression. Although atypical phenotype B cells dominated the response, we also identified a small subset of CD27+ classical Bmem establishing the co-existence of both pathways. Finally, our studies identify pre-switched B-cells as a reservoir for high-avidity expanded B cell clones. Collectively, our findings underscore the heterogeneity in donor-specific B cells, and provide a theoretic framework for targeting the resistance that must be overcome in the clinic if chronic humoral alloimmunity is to be controlled.

Similar findings of high affinity minimally mutated B cells were recently suggested in a pre-clinical allotransplantation model^15^ and also reported by others using lineage-tracing experiments in response to HIV antigens^14^. Importantly, unlike kidney transplantation where immunosuppression is usually minimized once the allograft fails, our study assessed the B-cell response in a transplant recipient maintained on triple-drug maintenance therapy which is necessary to mitigate the rejection process that can be fatal in heart transplantation. Hence, our study is unique in its focus on defining the donor-specific B cells escaping conventional immunosuppression that effectively constrains T-cell mediated rejection (as there was no evidence of TCMR on a paired biopsy). We speculate that these stochastically high-affinity B cells obviate the need for competitive and sustained T-cell help in the GC, and that lower levels of T-cell help may be sufficient to drive their differentiation into antibody-secreting cells in a GC-independent manner. In support of this model, mouse studies using SAP-knockout T cells have reported on the need for sustained B:T-cell interactions in the GC but not for extrafollicular (EF) B-cell responses^31,32^. Additional supportive evidence for alloreactive B-cell differentiation outside the GC comes from studies in which delaying CTLA-4Ig administration until day 7 post-sensitization suppressed the GC response but nonetheless permitted substantial early donor-specific IgG responses and memory B cell accumulation^33^.

We established that both germline and minimally mutated CD27^-^ B cells and affinity matured B cells with a classical CD27^+^ memory phenotype can both expand and co-exist in human transplantation. These findings raise important questions about the temporal development of humoral responses that require strong selection pressures to achieve a high avidity state versus those that arise stochastically with high avidity for alloantigen. One hypothesis is that affinity matured post-GC B cells develop during brief periods of CNI-reduction that preferentially allows for the development of T-follicular helper (Tfh) cells but not T-effector cells. Supporting this are the basic science observations of lower levels of TCR signaling promoting Tfh differentiation and higher TCR signals promoting T effector cells^34,35^. Low affinity B cells entering the GC compete for limited T-cell help to undergo affinity selection and emerge as affinity-matured Bmem or post-GC ASCs. One possibility is that affinity matured Bmem can also serve as antigen specific APCs to further support the persistence of cognate Tfh-like cells, which in turn, mediate extrafollicular (non-GC) Bmem differentiation into ASCs. An alternative possibility is that the differentiation of germline high affinity B cells into ASCs and Bmem requires low levels of extrafollicular T-cell help that develop to adequate levels despite maintenance immunosuppression. Subsequently this extrafollicular cognate T:B interaction generates primed Tfh-like cells that then mediate GC B-cell responses. Disentangling these possibilities will require additional studies in pre-clinical models, where temporal dynamics can be formally tested.

The finding that high avidity donor alloreactivity can be germline derived and differentiate into ASCs without the need for affinity maturation has important therapeutic implications. First, since B cells with high avidity germline BCRs may not rely on prolonged access to GC T-cell help, our findings provide a potential mechanism by which donor-reactive humoral immunity can develop and persist under T-cell centric immunosuppression. Furthermore, CNI-mediated partial suppression of TCR and/or IL-2 signals may skew T-cells towards a Tfh-like phenotype capable of providing help for T-dependent non-GC differentiation^12,35,36^. Second, germline B-cell alloreactivity may explain the limited efficacy of B-cell depleting therapies as newly emerging donor-reactive B cells receiving pre-existing T-cell help can rapidly differentiate into extrafollicular plasmablasts^37,38^. Consistent with this possibility, autoreactive B cells in rituximab-resistant immune thrombocytopenic purpura are derived from both the naïve and memory subsets^39^. Importantly, we and others have collectively put forth mouse, non-human primate, and human evidence that costimulation blockade more effectively suppresses humoral responses compared to CNI, by preventing early naïve T-cell priming and T-B crosstalk, decreasing the frequency of de novo DSA (including in high-risk HLA-DQ mismatches), and suppressing established humoral responses^19,20,22,24,40^. Belatacept is an already FDA licensed biologic that disrupts CD28-CD80/86 costimulation provides a readily available therapeutic strategy, although questions regarding dosing and optimal use in conjunction with other immunosuppressants remain to be answered. It is also possible that inhibiting the CD40/CD40L pathway which is critical for B-cell differentiation will be even more effective as suggested in a non-human primate xenotransplant model where treatment with an anti-CD40 monoclonal antibody provided long-term control of the humoral response^41^.

The single-case nature of this study is a limitation. However, the objective was not to define a uniform phenotype of chronic rejection but rather to establish the breadth of alloreactive B-cell heterogeneity that develops during chronic rejection despite contemporary immunosuppression. The prevalence of classical Bmem versus non-classical germline high affinity B cells in acute and chronic rejection in large cohort studies are urgently required. The immunobiology of donor HLA-DQ reactive B cells has not been well characterized, yet anti-HLA-DQ antibodies are common after transplant and strongly associated with pathogenicity^8,42^. This study also did not fully disentangle heterogeneity within the DN B-cell subset. Finally, it will be important to understand whether the signals required for donor-reactive B-cells to differentiate into ASCs are distinct for stochastically high-avidity versus low-avidity B cells. These questions are being addressed in ongoing work.

In conclusion, this study establishes the coexistence of affinity matured and germline derived high avidity donor HLA-DQ8 reactive B cells spanning pre- and post-switched subsets. The novel finding that stochastically generated, high avidity donor-reactive B cells are enriched for a non-canonical, antigen experienced phenotype that does not require GC-associated affinity maturation points to a mechanism of humoral escape under T-cell centric immunosuppression and transient B-cell depletion. These studies support the testable hypothesis that optimized use of costimulation blockade better constrains T-B cell crosstalk compared to conventional immunosuppression, and serve as a critical step towards preventing and reversing humoral-associated chronic rejection.

## Acknowledgements

The authors would like to thank Carol Benteljewski (University of Pittsburgh); Michael Kissner (Columbia University) and Elena Solomaha (University of Chicago, Biophysics Core (RRID:SCR_017915)). We also thank Dr. G. Kelsoe for providing the MS40L^low^ cell line, the NIH Tetramer Core Facility (contract number 75N93020D00005) for providing (HLA DQB1*03:02/DQA1*03:01 tetramers and monomers and DQB1*06:02/DQA1*01:02 monomers) and the NYU Langone Cytometry and Cell Sorting Laboratory (RRID: SCR_019179).

This work was in part supported by: National Institutes of Health U01 AI174997 (to MH), and R01AI142747, R01AI148705, P01AI097113 (to ASC).

## Data availability statement

The data that support the findings of this study are available from the corresponding author, [MH], upon reasonable request.

## Abbreviations

AMR: antibody mediated rejection
APC: antigen presenting cell
ASC: antibody secreting cell
BAFF: B-cell activating factor
BCR: B-cell receptor
Bmem: memory B cell
CAV: cardiac allograft vasculopathy
CNI: calcineurin inhibitor
DSA: donor specific antibody
EF: extrafollicular
GC: germinal center
HLA: human leukocyte antigen
IGHV: immunoglobulin heavy chain variable region gene
K_D_: equilibrium constant
MFI: mean fluorescence intensity
PBMC: peripheral blood mononuclear cell
PC: plasma cell
rAb: recombinant monoclonal antibody
SHM: somatic hypermutation
SPR: surface plasmon resonance
TCMR: T-cell mediated rejection
TCR: T cell receptor
Tfh: T-follicular helper

