## Supplementary material for "Donor HLA-DQ reactive B cells clonally expand under chronic immunosuppression and include atypical CD21^low^CD27^−^ B cells with high-avidity germline B-cell receptors": Figure S1

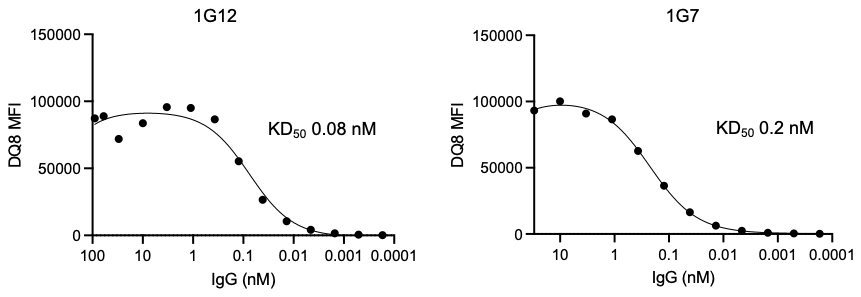

**Figure S1. Representative KD_50_ determined by dilution series. KD50 was defined as the concentration at 50% maximal MFI.**
