## Supplementary material for "Donor HLA-DQ reactive B cells clonally expand under chronic immunosuppression and include atypical CD21^low^CD27^−^ B cells with high-avidity germline B-cell receptors": Figure S2

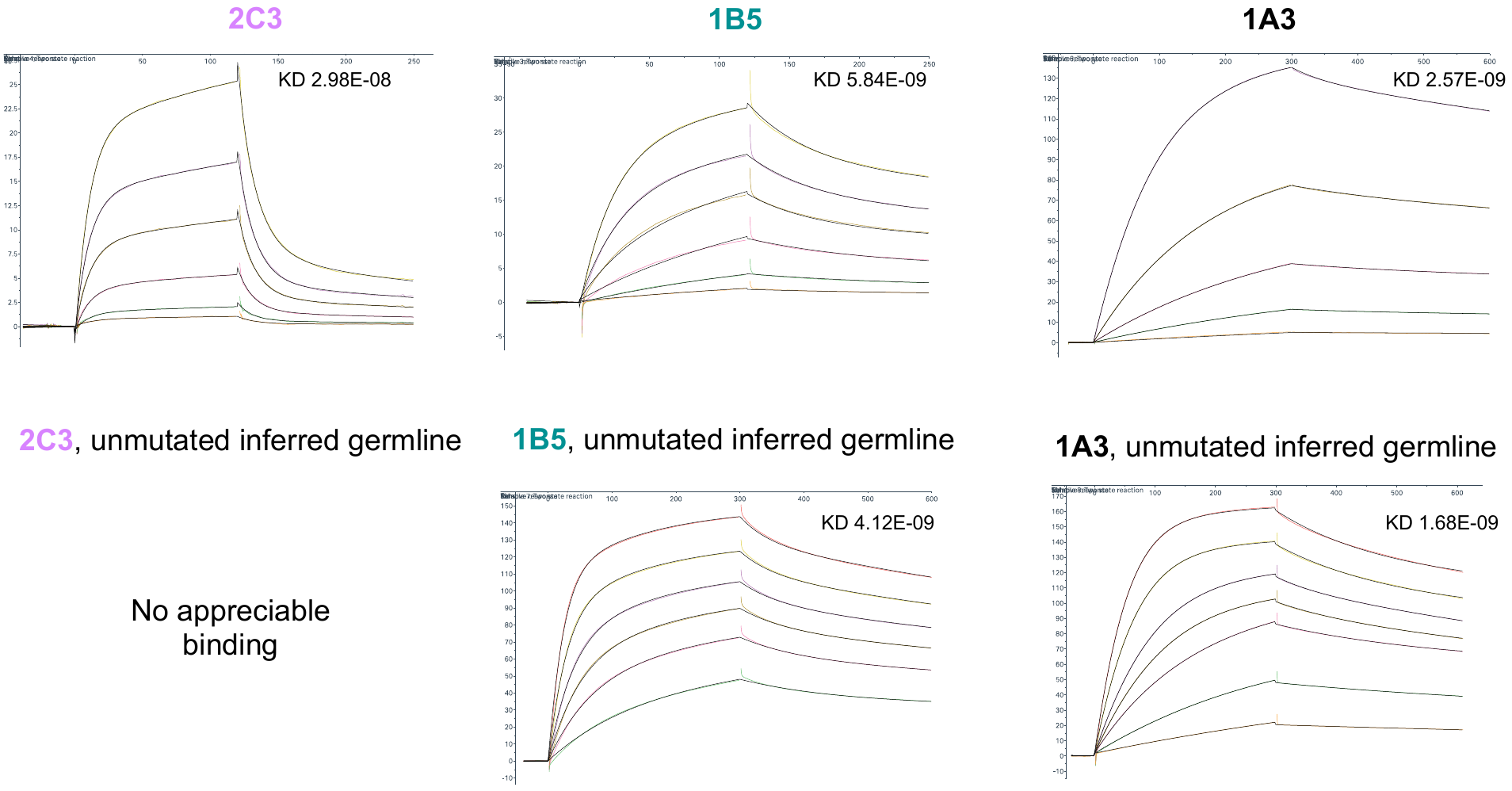

**Figure S2. SPR analysis of germline and reverted IgG1 rmAbs generated from donor reactive BCR heavy and light chain sequences.**
