## Supplementary figures and images for "Donor HLA-DQ reactive B cells clonally expand under chronic immunosuppression and include atypical CD21^low^CD27^−^ B cells with high-avidity germline B-cell receptors"

### Figure S3

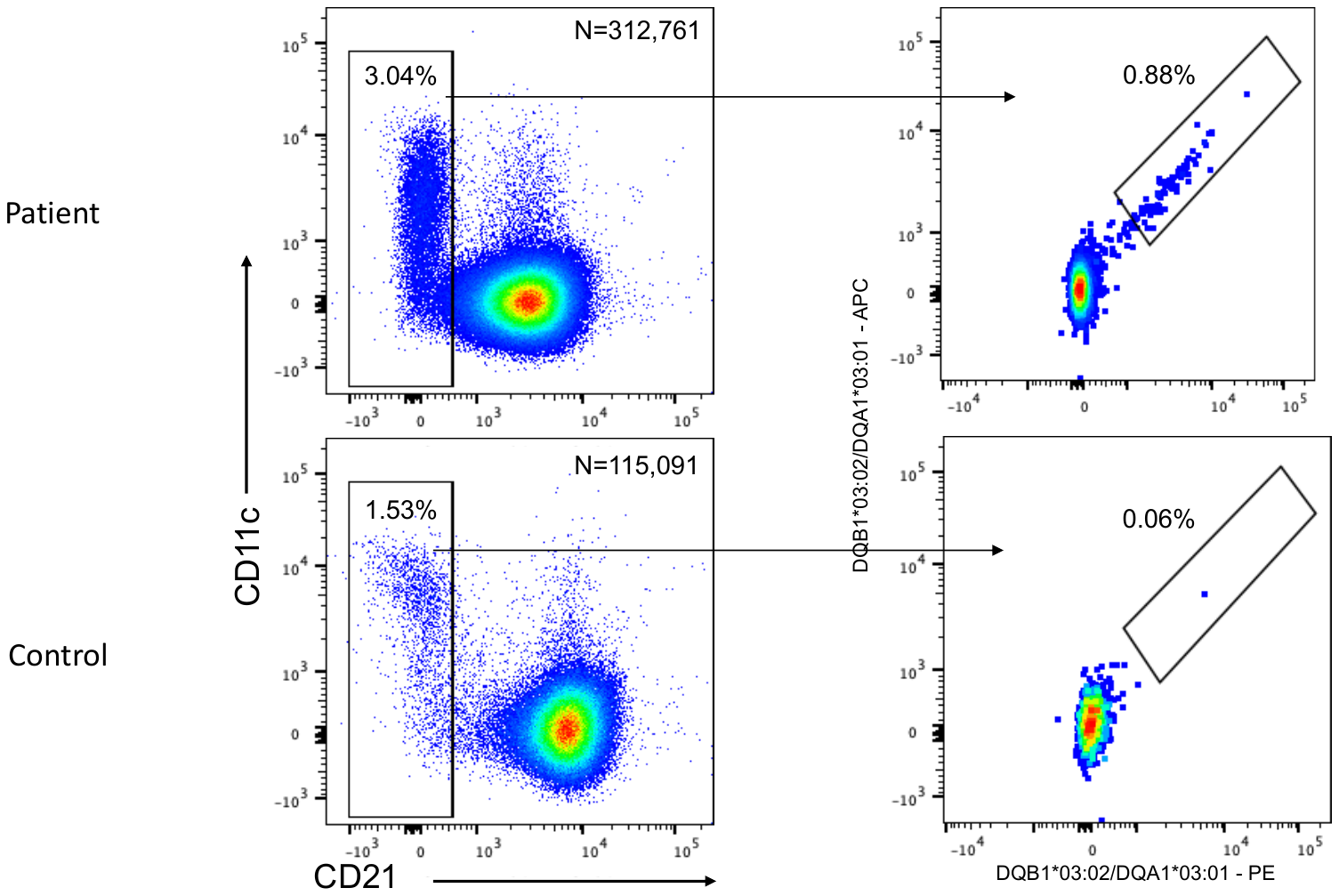

**Figure S3. Identification of a unique subset of IgD^+^CD27^-^ pre-switched donor reactive B cells.**
