## Supplementary material for "Donor HLA-DQ reactive B cells clonally expand under chronic immunosuppression and include atypical CD21^low^CD27^−^ B cells with high-avidity germline B-cell receptors": Figure S4

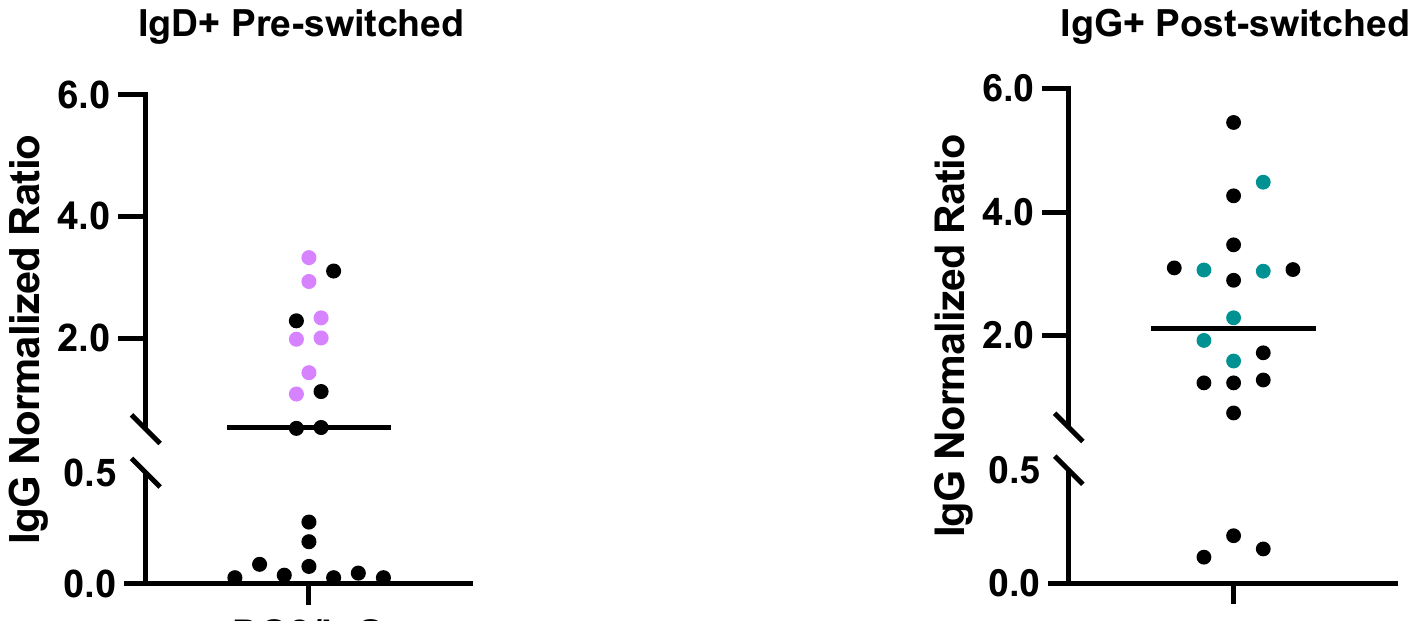

**Figure S4. IgG normalized HLA DQ8 MFI of pre-switched IgD+ (purple) and post-switched IgG+ (green) B-cell clones used for SPR analysis.**
