## Supplementary material for "Donor HLA-DQ reactive B cells clonally expand under chronic immunosuppression and include atypical CD21^low^CD27^−^ B cells with high-avidity germline B-cell receptors": Figure S5

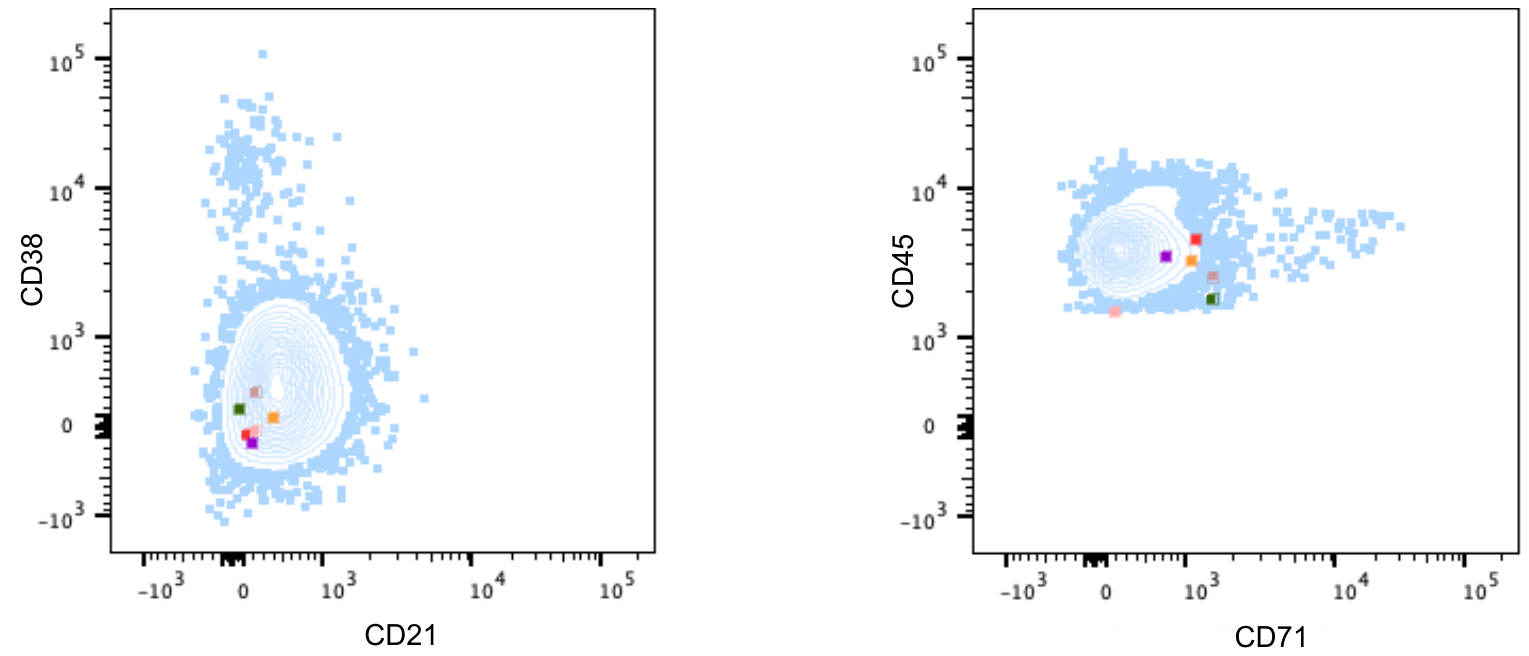

**Figure S5. Phenotype of donor reactive B-cell clones that persisted after treatment with rituximab.**
