## Supplementary material for "Donor HLA-DQ reactive B cells clonally expand under chronic immunosuppression and include atypical CD21^low^CD27^−^ B cells with high-avidity germline B-cell receptors": Table S1

**Table S1. Resources and Reagents**

| **REAGENT** | **SOURCE** | **IDENTIFIER** |
| --- | --- | --- |
| **Antibodies** | | |
| BV421 Mouse Anti-Human CD21 (clone B-ly4) | BD Bioscience | 562966 |
| BV605 Mouse Anti-Human CD11c (clone B-ly6) | BD Bioscience | 563929 |
| PE/Cyanine5 Mouse anti-human CD3 (clone HIT3a) | BioLegend | 300310 |
| PE/Cyanine5 Mouse anti-human CD14 (clone M5E2) | BioLegend | 301864 |
| FITC Mouse Anti-Human CD19 (clone HIB19) | Thermo Fisher Scientific | 11-0199-42 |
| BV510 Mouse Anti-Human IgD (clone IA6-2) | BioLegend | 348219 |
| BV650 Mouse Anti-Human CD71 (clone CY1G4) | BioLegend | 334116 |
| AF700 Mouse Anti-Human CD20 (clone 2H7) | BioLegend | 302322 |
| Spark UV 387 Mouse Anti-Human CD45 (clone HI30) | BioLegend | 304085 |
| PE/Dazzle 594 Mouse Anti-Human CD27 (clone M-T271) | BioLegend | 356421 |
| APC/Cyanine7 Mouse Anti-Human CD38 (clone HB7) | BioLegend | 356616 |
| BV786 Mouse Anti-Human IgG (clone G18-145) | BD Bioscience | 564230 |
| Biotin-conjugated Goat Anti-Human IgG | SouthernBiotech | 2040-08 |
| PE-conjugated Goat Anti-Human IgG | SouthernBiotech | 2040-09 |
| ChromPure Human IgG, whole molecule | Jackson ImmunoResearch | 009-000-003 |
| **Recombinant proteins, monomers, tetramers and other reagents** | | |
| Recombinant human IL-2 | Peprotech | 200-02 |
| Recombinant human IL-21 | Peprotech | 200-21 |
| Recombinant human IL-4 | Peprotech | 200-04 |
| Recombinant human BAFF | Peprotech | 310-13 |
| Invitrogen UltraComp eBeads | Thermo Fisher Scientific | 50-112-9040 |
| FCS HyClone (Fetal Calf Serum) | Thermo Fisher Scientific | SH3007003.03 |
| Carboxyl Blue Particle Array Kit, Even # peaks | Spherotech | CPAK-5067-5B |
| Propidium Iodide | Biolegend | 421301 |
| Benzonase Nuclease | Sigma-Aldrich | e8263 |
| Recombinant IgG1, Lot# KH18JA28007 | Sino Biological | NYU4-1 |
| Recombinant IgG1, Lot# KH18JA28008 | Sino Biological | NYU4-2 |
| Recombinant IgG1, Lot# KH18JA28009 | Sino Biological | NYU4-3 |
| Recombinant IgG1, Lot# KH18JA28010 | Sino Biological | NYU4-4 |
| Recombinant IgG1, Lot# KH18JA28011 | Sino Biological | NYU4-5 |
| Recombinant IgG1, Lot# KH18JA28012 | Sino Biological | NYU4-6 |
| DQB1*03:02/DQA1*03:01 APC-labeled Tetramer | NIH Tetramer Core | N/A |
| DQB1*03:02/DQA1*03:01 PE-labeled Tetramer | NIH Tetramer Core | N/A |
| DQB1*06:02/DQA1*01:02 Biotinylated Monomer | NIH Tetramer Core | N/A |
| SARS-CoV-2 (2019-nCoV) Spike S1-His Recombinant Protein, Biotinylated | Sino Biological | 40591-V08H-B |
| **Cell lines** | | |
| MS40L^low^ | Gift of Dr. G. Kelsoe | N/A |
| **Sequencing and recombinant antibodies** | | |
| B-cell receptor sequencing: iRepertoire (Huntsville, AL) | | |
| Recombinant antibodies: Sino Biological | | |
| **Software and algorithms** | | |
| FlowJo | Treestar Inc | https://www.flowjo.com/ |
| FACSDIVA | BD Biosciences | http://www.bdbiosciences.com/us/instruments/research/software/flow-cytometry-acquisition/bd-facsdiva-software/m/111112/overview |
| IMGT/V-QUEST | Brochet et al., 2008 | http://www.imgt.org/IMGT_vquest/vquest |
| Prism 10.1.1 | GraphPad | https://www.graphpad.com/ |
