## Supplementary material for "Donor HLA-DQ reactive B cells clonally expand under chronic immunosuppression and include atypical CD21^low^CD27^−^ B cells with high-avidity germline B-cell receptors": Table S2

**Table S2. IGHV somatic hypermutation characteristics for switched IgG+ donor-reactive B cells**

| **Clone** | **Number of mutations** | **Silent** | **Non-silent** | **Non-silent/**  **silent** | **Mutation**  **frequency** |
| --- | --- | --- | --- | --- | --- |
| 1A7 | 23 | 8 | 15 | 1.88 | 7.85% |
| 2C3 | 17 | 3 | 14 | 4.67 | 5.69% |
| 1F5 | 15 | 7 | 8 | 1.14 | 5.12% |
| 2C11 | 13 | 5 | 8 | 1.60 | 4.35% |
| 1A2 | 13 | 2 | 11 | 5.50 | 4.38% |
| 1A5 | 11 | 2 | 9 | 4.50 | 3.74% |
| 1A3 | 7 | 3 | 4 | 1.33 | 2.36% |
| 2A9 | 5 | 3 | 2 | 0.67 | 1.70% |
| 2A1 | 5 | 1 | 4 | 4.00 | 1.70% |
| 1B5 | 5 | 1 | 4 | 4.00 | 1.70% |
| 2A7 | 5 | 3 | 2 | 0.67 | 1.77% |
| 1H12 | 3 | 2 | 1 | 0.50 | 1.01% |
| 1A4 | 2 | 1 | 1 | 1.00 | 0.67% |
| 1E9 | 1 | 0 | 1 | 0.00 | 0.34% |
| 2B5 | 1 | 0 | 1 | 0.00 | 0.35% |
| 1A6 | 1 | 0 | 1 | 0.00 | 0.34% |
| 1A10 | 0 | 0 | 0 | 0.00 | 0.00% |
| 1G12 | 0 | 0 | 0 | 0.00 | 0.00% |
| F12 | 0 | 0 | 0 | 0.00 | 0.00% |
| **Average** | **6.68** | **2.15** | **4.52** | **1.65** | **2.27%** |
